# Myogenesis defects in a patient-derived iPSC model of hereditary GNE myopathy

**DOI:** 10.1101/2021.01.04.425299

**Authors:** Rebecca E Schmitt, Douglas Y Smith, Dong Seong Cho, Lindsey A Kirkeby, Zachary T Resch, Teerin Liewluck, Zhiyv Niu, Margherita Milone, Jason D Doles

## Abstract

Hereditary muscle diseases are disabling disorders lacking effective treatments. UDP-N-acetylglucosamine-2-epimerase/N-acetylmannosamine kinase (GNE) myopathy is an autosomal recessive distal myopathy with rimmed vacuoles that typically manifests in late adolescence/early adulthood. GNE encodes an enzyme that is the rate-limiting step in sialic acid biosynthesis which is necessary for proper function of numerous biological processes. Outside of the causative gene, very little is known about the mechanisms contributing to the development of GNE myopathy. In the present study we aimed to address this knowledge gap by querying underlying mechanisms of GNE myopathy using a patient-derived induced pluripotent stem cell (iPSC) model. Muscle and skin biopsies were acquired from two patients with GNE myopathy that presented with distinct histopathological features. Control and patient-specific iPSCs were derived from skin fibroblasts and differentiated down a skeletal muscle lineage in a three-stage process analogous to muscle development and muscle regeneration. Initial studies revealed: 1) the ability of patient-derived GNE iPSC clones to recapitulate key characteristics of the human pathology including TDP-43 accumulation and evidence of dysregulated autophagy, and 2) a striking defect in myogenic progression of the more severe GNE iPSC clone. Single-cell RNA sequencing time course studies were then performed to more rigorously explore myogenesis defects. Cluster-based bioinformatics analyses revealed clear differences between control and GNE iPSC-derived muscle precursor cells (iMPCs). On a transcriptional level, late stage GNE iMPCs resembled that of early stage control iMPCs, confirming stalled myogenic progression on a molecular level. Comparative expression and pathway studies revealed EIF2 signaling as a top signaling pathway altered in GNE iMPCs. In summary, we report a novel *in vitro*, iPSC-based model of GNE myopathy and implicate defective myogenesis as a likely novel contributing mechanism to the etiology of GNE myopathy.

**SUMMARY STATEMENT:** Development of a novel cell-based model of GNE myopathy, utilizing GNE patient-derived samples, which recapitulates human disease characteristics, uncovered myogenic differentiation defects, and can elucidate possible mechanistic contributors to the disease.

## INTRODUCTION

UDP-N-acetylglucosamine-2-epimerase/N-acetylmannosamine kinase (GNE) myopathy is an early adulthood onset, autosomal recessive, rare myopathy with a prevalence of ~1-9/1,000,000 (Carrillo et al., 2018). Some communities, such as the Persian Jewish community, have a much higher prevalence of ~1/1,500 (Salama et al., 2005). Historically, prior the identification of the underlying genetic defect, GNE myopathy was independently described by various investigators as Nonaka distal myopathy, distal myopathy with rimmed vacuoles, vacuolar myopathy sparing quadriceps, and inclusion body myopathy 2 (IBM2) (Nonaka et al., 1981; Argov and Yarom, 1984; MitraniRosenbaum et al., 1996; Huizing et al., 2014). On a molecular level, the *GNE* gene encodes a protein containing two enzymatic domains, the epimerase and kinase domain, which are involved in sequential steps of the biosynthesis of sialic acid. Sialic acids are a common terminal sugar on glycolipids and glycoproteins, and are involved in diverse biological functions from aging to immune responses (Schauer, 2009). To date more than 200 mutations have been found in patients suffering from GNE myopathy and these mutations can be either homozygous, as seen in specific ethnic populations, or compound heterozygous, and occurring in either the same or different domains of GNE (Carrillo et al., 2018). Clinically, GNE myopathy manifests with distal lower limb weakness resulting in foot drop. Over time, the muscle weakness spreads to proximal muscles and eventually to the upper limb muscles but classically spares the quadriceps. Within two decades from disease onset, patients frequently have impaired mobility leading to a need for wheelchair assistance (Argov and Yarom, 1984; Pogoryelova et al., 2018). Pathologically, GNE myopathy is characterized by rimmed vacuoles, congophilic inclusions, protein aggregates, and enhanced lysosomal activity as evidenced by increased acid phosphatase reactivity (Zhang et al., 2018; Pogoryelova et al., 2018; Carrillo et al., 2018).

There are currently no effective treatments for GNE myopathy. Numerous organismal GNE models have been developed to address this issue, however most of these models have significant flaws that limit their ability to provide insights into disease mechanism or treatment. Several are briefly discussed below. A significant concern is that many murine GNE models simply do not develop any muscle abnormalities or myopathic features typical of the human disease. For example, some GNE models feature glomerulopathy, which is not found in GNE patients (Galeano et al., 2007; Kakani et al., 2012; Ito et al., 2012). Concerns aside, there is one mouse model combining a GNE null allele with transgenic expression of a human GNE D176V allele that appears to rise above the rest. This mouse develops clinical characteristics comparable to those observed in GNE patients (Malicdan et al., 2007). For example, rimmed vacuoles, increased acid phosphatase staining, and increased LAMP1 and LAMP2 staining are evident in cross-sectional analyses of the gastrocnemius muscle. Furthermore, these mice display loss of motor strength after the age of 30 weeks. As GNE null mice are embryonic lethal (Schwarzkopf et al., 2002; Galeano et al., 2007), maintaining this model and generating experimental mice is challenging as the required mating scheme results in only 9% of the pups having the genotype of interest (Malicdan et al., 2007). Thus, while potentially useful for studying disease etiology, this model is likely ill-suited for drug screening and/or medium- to large-scale pre-clinical intervention studies. In addition to murine models, GNE myopathy can also be modeled using zebrafish (Daya et al., 2014). Human and zebrafish GNE share the same functional domains and have a 90% similarity. In one study, morpholino-mediated GNE knockdown resulted in variable morphological deformities ranging from relatively normal to severe in regards to overall size and tail/trunk development. Additionally, skeletal muscle myofibers were found to be highly disorganized, an observation likely to explain the decreased larval locomotor activity in the normal/less severely affected GNE knockdown larvae compared to controls (Daya et al., 2014). Unfortunately, despite the promise of these preliminary observations, little has since been pursued using this model, particularly with respect to mechanisms downstream of GNE that could contribute to these muscle phenotypes or in the context of drug screening, one strength of the zebrafish system.

The role of mutated GNE in the development of muscle weakness and human myopathology are unknown. A number of past and current clinical trials using sialic acid, or its precursors, to treat GNE myopathy have yielded underwhelming results (Lochmuller et al., 2019). These results raise the likely probability that GNE has additional functions, beyond its role in sialic acid metabolism, that are relevant to disease development and progression. Revealing novel mechanistic insights will require a model that reliably recapitulates the human phenotype, is feasible to obtain, and is cost-effective. In this study, we report development of an *in vitro GNE* myopathy model using patient-derived induced pluripotent stem cells (iPSCs). Utilizing this model, we identify previously unrecognized deficiencies in myogenesis. We corroborate these observations using longitudinal single-cell RNA sequencing analyses. Finally, we identify GNE-associated signaling pathway alterations, revealing potentially novel therapeutic opportunities to improve GNE myopathy pathophysiology.

## RESULTS

### Histopathological evaluation of GNE patient muscle biopsies

The first goal of this study was to evaluate the histopathological severity of two GNE patient biopsies by assessing standard GNE myopathy characteristics (Carrillo et al., 2018; Nishino et al., 2015) in order to establish a baseline for eventual comparison to patient-derived cells. First, hematoxylin and eosin (H&E) staining revealed that both GNE patient samples, GNE^1^ and GNE^2^, contained myofibers of variable size including atrophic fibers (Fig. 1A-B, asterisk). Second, both samples featured rimmed vacuoles, which were more abundant in GNE^1^ compared to GNE^2^ (Fig. 1A-B, arrows). Third, Congo Red staining revealed congophilic inclusions (Fig. 1 C-D, arrows) while acid phosphatase staining showed patchy over reactivity especially in the vacuoles (Fig. 1E-F, arrows). Fourth, autophagy pathway activity was queried by staining muscle cross-sections for p62 reactivity (Korolchuk et al., 2009; Liu et al., 2016). Here, two distinct patterns of p62 were observed. The first pattern observed was focal accumulation of p62 (arrows) while the second pattern was diffuse p62 staining throughout the myofiber (asterisk) (Fig. 1G-H). Both mutation-confirmed GNE patient muscle biopsies exhibited the histopathological GNE hallmarks, with GNE^1^ qualitatively exhibiting a more severe pathological phenotype.

**Figure 1.**
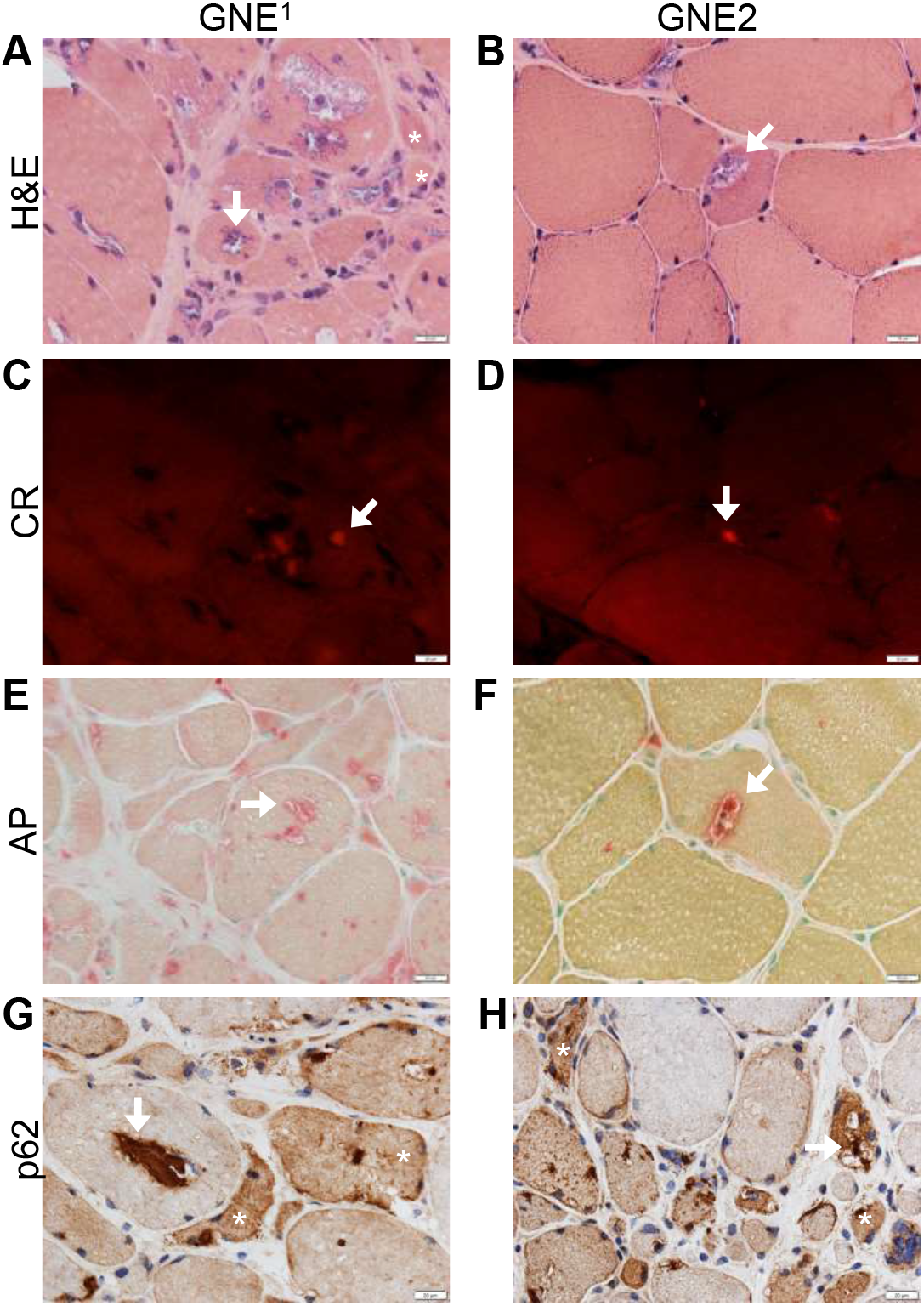
Characterization of GNE patient muscle biopsies. (A-B) Hematoxylin and eosin (H&E) stained tissue cross-sections of two patients with GNE myopathy, GNE^1^ (left) and GNE^2^ (right), exhibit rimmed vacuoles (arrows) and muscle fiber size variability, including presence of atrophic fibers (asterisk). (C-D) Congo Red stained sections highlighting the presence of congophilic inclusions (arrows). (E-F) Acid phosphatase over-reactivity within vacuoles (arrows) and in a punctuate fashion outside the vacuoles, suggesting lysosomal dysfunction. (G-H) Ectopic p62 immunoreactivity occurs focally (arrow) or diffusely (asterisk) in numerous muscle fibers. All images are 40X, scale bars: 20 μm.

### Patient-derived induced pluripotent stem cells (iPSCs) exhibit GNE myopathy hallmarks and differentiate poorly into mature skeletal muscle myotubes

Punch biopsies from both patients were collected and fibroblast cultures established. Healthy (no myopathy) control and GNE patient-derived fibroblasts were reprogrammed into induced pluripotent stem cells (iPSCs) using Sendai virus-Cytotune 2.0. Two clones were derived from the healthy control line and three clones derived from each of the two GNE patient lines (Table 1). All clones underwent rigorous quality control characterization routinely performed in the iPSC field. In brief, iPSC identity was confirmed by short tandem repeat (STR) analysis, found negative for mycoplasma, and exhibited normal karyotypes via G-banding [data not shown (DNS)]. Pluripotency markers were assessed and all clones were positive for Oct4, SSEA, Nanog, and TRA-1-60 (Fig. 2A and DNS) with no gross abnormalities apparent. Since it has been shown that the iPSC reprogramming process can itself introduce genetic variation/drift (Liang and Zhang, 2013), we utilized Sanger Sequencing to confirm the retention of GNE mutations post-reprogramming in one set of clones from each line. The control line demonstrated homozygous genotypes containing wild type alleles, seen as the presence of a single nucleotide peak (Fig. 2Bi-ii and Di-ii, red boxes). Conversely, GNE^1^ contained double peaks of relatively equal size for both G and A at c.479 and c.2179 (Fig. 2Ci-ii, red boxes) indicating the presence of a single mutation in both the epimerase and kinase domains. GNE^2^ contained double peaks for G and T at GNE c.1839 and G and A at c.2218, respectively (Fig. 2Ei-ii, red boxes), which corresponds to two mutations in the kinase domain of GNE. These Sanger Sequencing results matched the original patient compound heterozygous mutations found at diagnosis (Table 1). Lastly, to confirm germline differentiation capacity, all clones were tested for proficiency to differentiate into all three germ layers. Standard ectoderm, endoderm, and mesoderm markers were qualitatively assessed post-differentiation (Fig. 2F-H). All control and GNE patient-derived iPSC clones were capable of expressing Nestin and Pax-6 post-ectoderm differentiation (Fig. 2F and DNS), FoxA2 and Sox17 after endoderm differentiation (Fig. 2G and DNS), and CD31 and NCAM for cells differentiated down a mesoderm lineage (Fig. 2H and DNS). These results demonstrate successful reprogramming of patient-derived fibroblasts into functional iPSCs.

**Table 1.**
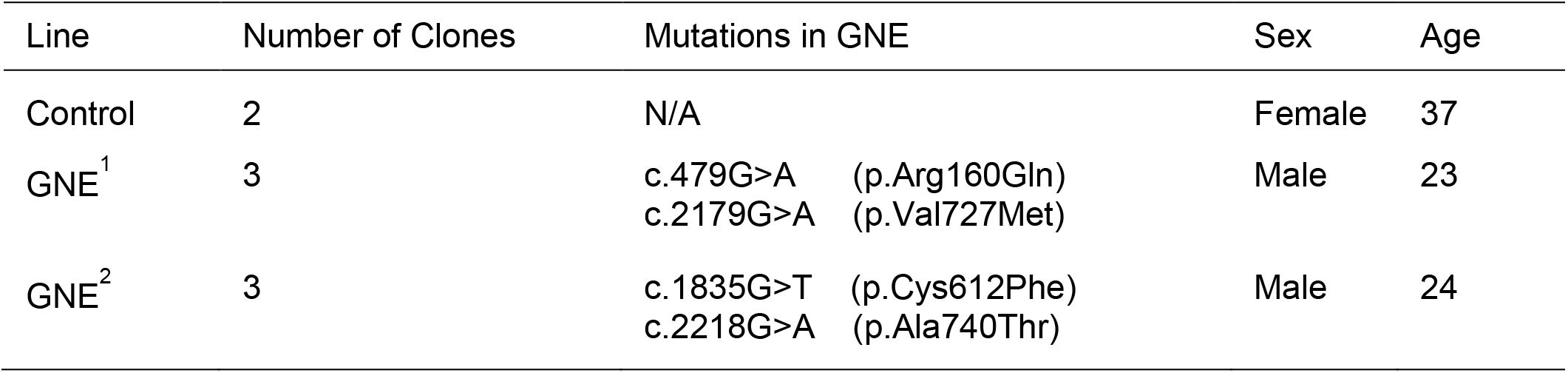
Features of healthy or GNE patients of which iPSC lines were derived. Control, GNE^1^, and GNE^2^ are listed by name and subsequent number of clones per line, presence of GNE mutations, sex, and age are described. N/A is not applicable.

| Line | Number of Clones | Mutations in GNE | Sex | Age |
| --- | --- | --- | --- | --- |
| Control | 2 | N/A | Female | 37 |
| GNE<sup>1</sup> | 3 | c.479G>A (p.Arg160Gln)<br>c.2179G>A (p.Val727Met) | Male | 23 |
| GNE<sup>2</sup> | 3 | c.1835G>T (p.Cys612Phe)<br>c.2218G>A (p.Ala740Thr) | Male | 24 |

**Figure 2.**
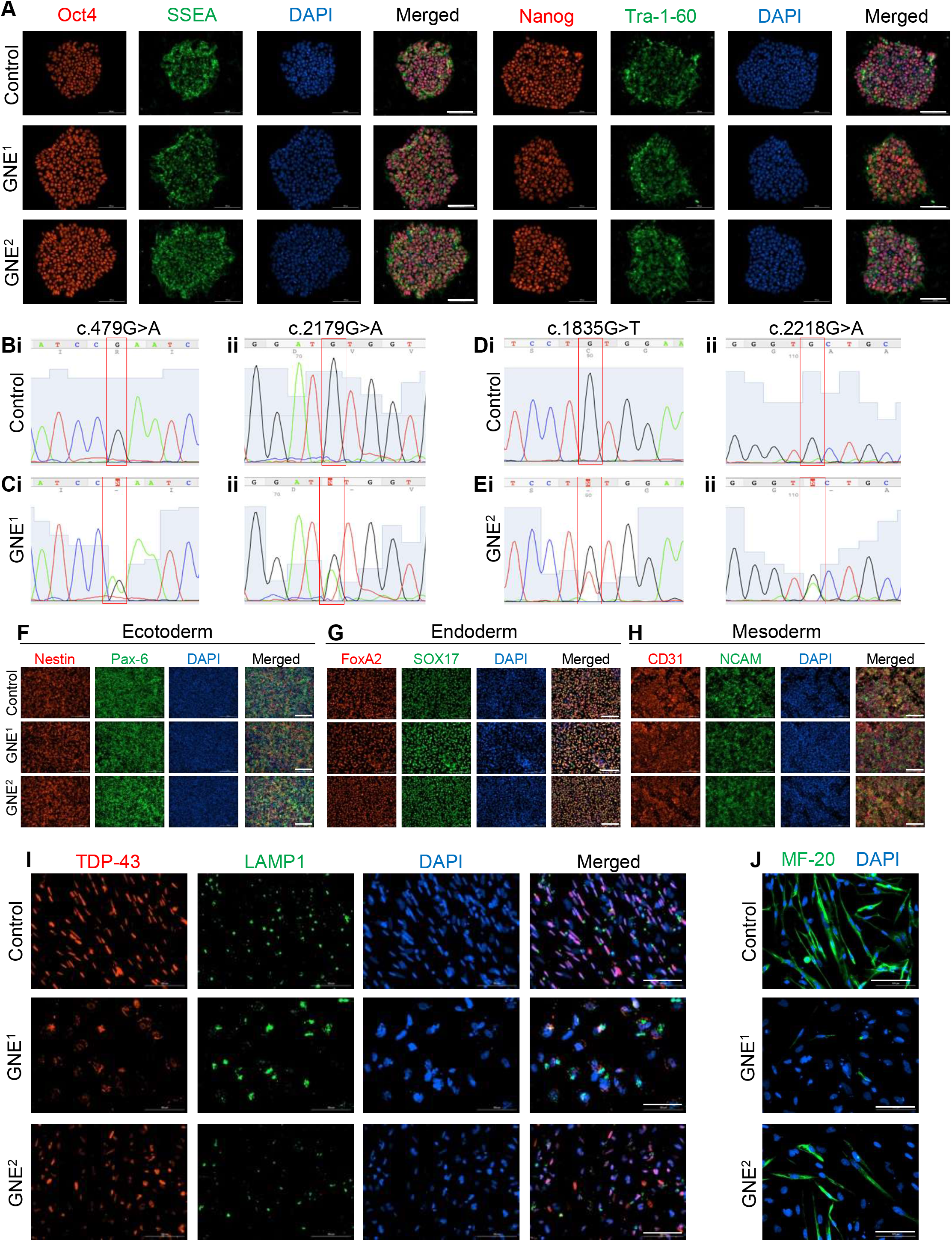
Functional iPSC reprogramming and characterization. (A) Control (top), GNE^1^ (middle), and GNE^2^ (bottom) iPSCs were assessed for expression of standard pluripotency markers by immunofluorescence (IF). All clones expressed Oct4, SSEA, Nanog, and Tra-1-60. (B-D) Sanger sequencing plots confirming the presence of the compound heterozygous patient mutations. Control samples contain the wild type alleles at all mutant locations (red boxes, Bi-ii and Di-ii), whereas GNE^1^ are heterozygous for wild type and mutant alleles (red boxes, Ci-ii) as is GNE^2^ (red boxes, Ei-ii). (F-H) IF images highlighting germline differentiation capacity. Differentiated iPSCs were capable of expressing standard ectoderm (Nestin and Pax-6) (F), endoderm (FoxA2 and SOX17) (G), and mesoderm (CD31 and NCAM) markers (H). (I-J) IF images depicting GNE marker and myosin heavy chain expression. Control, GNE^1^, and GNE^2^ iPSCs were differentiated to myotubes and stained for (I) standard GNE markers including TDP43 (red), LAMP1 (green), and DAPI (I) or myosin heavy chain (MF-20, green) and DAPI (blue) (J). All images are representative and taken at 20X, scale bars: 100 μm.

Next, we sought to determine the extent to which GNE patient-derived cells exhibited characteristics of GNE myopathy *in vitro*. Expression patterns of TAR DNA Binding Protein 43 (TDP-43) and the Lysosomal Associated Membrane Protein 1 (LAMP1) were assessed in control and GNE patient-derived cells subjected to a 3-step myogenic differentiation protocol to form skeletal muscle myotubes. TDP-43 is known to be present in the protein aggregates of GNE myopathy (Kusters et al., 2009) as well as having ectopic sarcoplasmic expression in other myopathies with rimmed vacuoles, including sporadic inclusion body myositis (Sandell et al., 2016; Nalbandian et al., 2011; Salajegheh et al., 2009). In control derived myotubes, TDP-43 signal co-localized with the nuclear stain, 4’,6-diamidino-2-phenylindole (DAPI), as expected (Fig. 2I). Strikingly, GNE^1^ myotubes exhibited a strong TDP-43 signal in some cytoplasmic regions (Fig. 2I), with little normal TDP-43 and DAPI overlap. GNE^1^ also exhibited large LAMP1 puncta, a lysosome l marker indicating the presence of autophagy (Eskelinen, 2006) (Fig. 2I). Alternatively, GNE^2^ myotubes appeared to have an intermediate phenotype between control and GNE^1^ in regards to expression patterns of TDP-43 and LAMP1 (Fig. 2I). Finally, compared to controls, we observed almost no MYH1 (using the monoclonal antibody MF-20) reactivity in GNE^1^ and reduced expression in GNE^2^, suggesting that myogenic differentiation is impaired in GNE myopathy iPSCs progressing down a skeletal muscle lineage (Fig. 2J).

### Patient-derived induced pluripotent stem cells (iPSCs) exhibit alterations in myogenic regulatory factor (MRF) expression during myogenic differentiation

Given the striking inability of GNE^1^ iPSCs to form mature skeletal muscle myotubes, we next sought to query stage-specific myogenic regulatory factor (MRF) protein expression in differentiating control and GNE iPSCs (hereby referred to as induced myogenic progenitor cells, or iMPCs). Three iMPC stages were assessed that correspond to early, intermediate and late stages of myogenesis: 1) stage 1-satellite-like cells, 2) stage 2-myoblasts, and 3) stage 3-myocytes/myotubes (Fig. 3A, green diamonds). Immunofluorescence was performed on two clones from the control patient-derived line and three clones from both GNE myopathy patient-derived lines. Nuclear-overlapping MRF expression was quantified as a percentage of total nuclei. At stage 1, both GNE^1^ and GNE^2^-derived samples exhibited a statistically significant decrease (18.85% ± 6.84% and 17.84% ± 5.92%, respectively) of Pax3 percent positive nuclei when compared to controls, while no difference was observed in Myf5 positive nuclei (Fig. 3B-C). GNE^1^ displayed an increased fraction of MyoD positive nuclei compared to control (24.84% ± 4.68%), whereas GNE^2^ did not (Fig. 3D-E). No differences in Myf5, MyoD, or MyoG positive nuclei were observed in stage 2 cells (Fig. 3F-I). A statistically significant reduction in the fraction of Pax3 positive nuclei persisted in both GNE^1^ and GNE^2^ samples at stage 2 compared to controls (44.78% ± 8.77% and 33.42% ± 8.77%, respectively) (Fig. 3H-I). At stage 3, MF20 was abundantly detected in control myotubes, but very little to no expression was found in the iMPC sample derived from GNE^1^ iPSCs (Fig. 3J). The number of myofibers per high power field was calculated and both GNE^1^ and GNE^2^ samples had decreased myofiber counts compared to control (27.27 ± 6.3 and 21.07 ± 6.3, respectively) (Fig. 3K). In addition, MyoG extensively overlapped with DAPI in control myotubes, but was rarely detected as an overlap in GNE^1^ and was significantly increased in control cells compared to both GNE^1^ and GNE^2^ (34.88% ± 8.3% and 34.4% ± 8.3%, respectively) (Fig. 3J and L).

**Figure 3.**
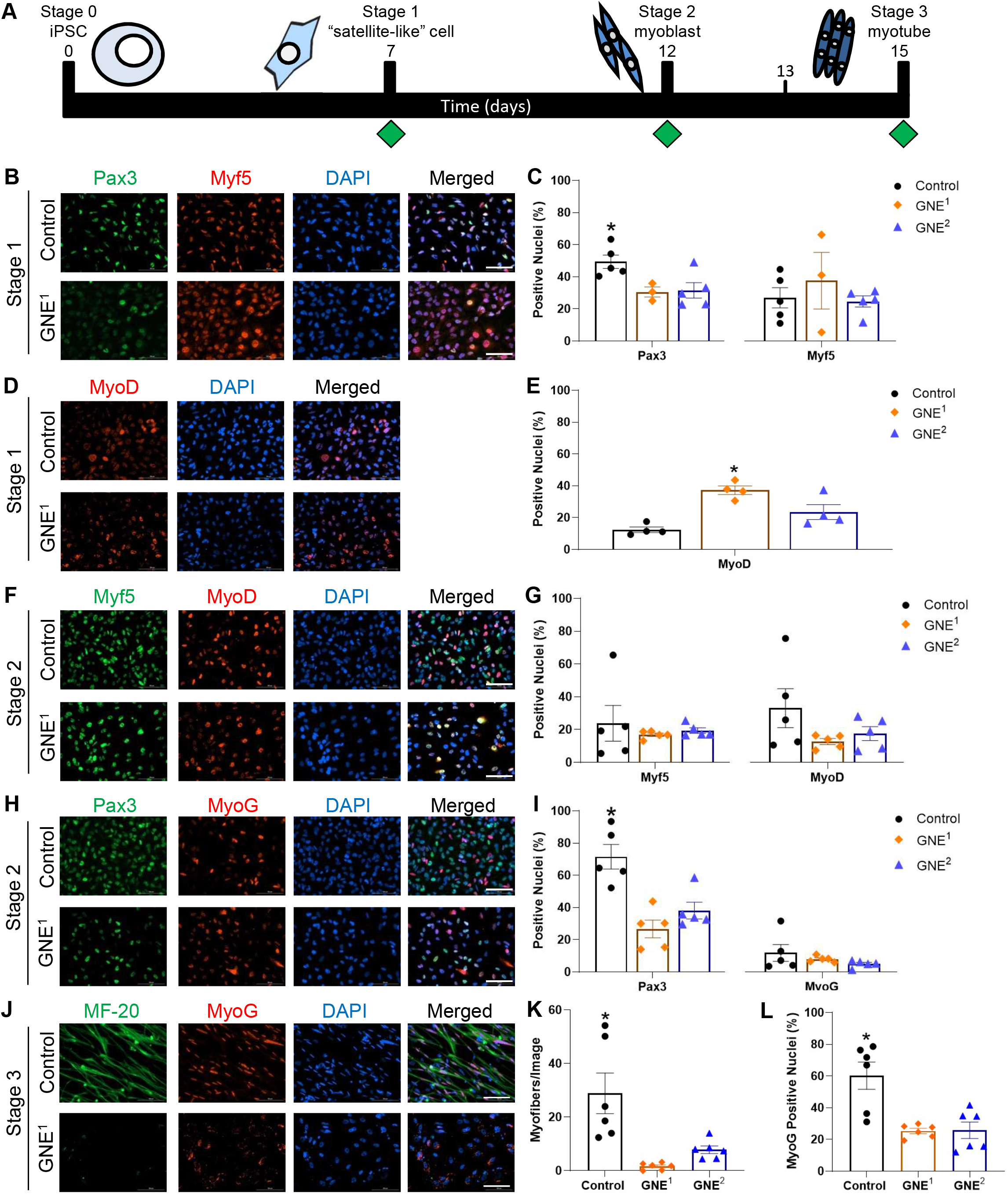
Muscle regulatory factor (MRF) expression during myogenic progression of healthy and GNE patient-derived iPSCs. (A) A schematic depicting the experimental timeline. Green diamonds represent the time of sample preparation for immunofluorescence. (B-E) Satellite-like cell, or stage 1 iMPCs. MRF protein expression in Control (top) and GNE^1^ (bottom) cells stained with (B) Pax3 (green), Myf5 (red), and DAPI (blue) or (D) MyoD (red) and DAPI (blue). (C, E) Quantification of the percent positive nuclei from control (black), GNE^1^ (orange), and GNE^2^ (blue) in images B and D, respectively. (F-I) Protein expression of myogenic transcription factors of stage 2 iMPCs. (F, H) Control (top) and GNE^1^ (bottom) stage 1 cells stained with (F) Myf5 (green), MyoD (red), and DAPI (blue) and (H) Pax3 (green), MyoG (red), and DAPI (blue). (G, I) Quantification of the percent positive nuclei from control (black), GNE^1^ (orange), and GNE^2^ (blue) in images F and H, respectively. (J-L) Control (top) and GNE^1^ (bottom) iMPCs were differentiated to myotubes (stage 3) and stained for myogenic markers MyoG (red) and myosin heavy chain (green). (K) Quantification of number of myofibers per 20X image. (L) Quantification of the percent positive nuclei from control (black), GNE^1^ (orange), and GNE^2^ (blue). All images are 20X representative images, scale bar: 100 μm. All statistical analyses were performed using one-way ANOVA with Bonferroni Multiple Comparisons (BMCs); *p<0.05.

### Widespread global transcriptomic differences between healthy and GNE patient-derived iPSC/iMPC samples during myogenic differentiation

Next, we sought to query molecular (transcriptional) alterations in differentiating control and GNE patient iPSCs/iMPCs. We selected three time points, iPSC (time point 0), satellite-like cells or early iMPCs (time point 1), and shortly after myoblast formation but not yet myotubes, or late iMPCs (time point 2) for these studies (Fig. 4A, red stars). Single cells were prepared using a 10x Genomics workflow and sequenced on an Illumina HiSeq4000 platform. Quality control (QC) was performed using Illumina software and the Seurat package in R (Table S1). Six samples in total were analyzed containing one clone from the control and one clone from GNE^1^ (these specific clones are depicted in Figs. 2 and 3). Control samples at time point 0 (Control-TP0), 1 (Control-TP1), and 2 (Control-TP2) were combined and visualized by the Uniform Manifold Approximation and Projection (UMAP) (Figure 4B). The control samples demonstrate distinct clustering of each cell stage, indicating relatively dissimilar transcriptomic profiles across differentiation, as expected. GNE^1^ cells at time points 0 (GNE-TP0), 1 (GNE-TP1), and 2 (GNE-TP2) were similarly visualized by UMAP and exhibited distinct stage-specific clustering (Fig. 4C). We combined control and GNE samples in a single UMAP plot and observed that control and GNE samples clustered reasonably well together at TP0 and TP1. Strikingly, GNE-TP2 iMPCs clustered just outside of control/GNE-TP1 samples, whereas Control-TP2 clustered distantly in the lower left quadrant (Fig. 4D).

**Figure 4.**
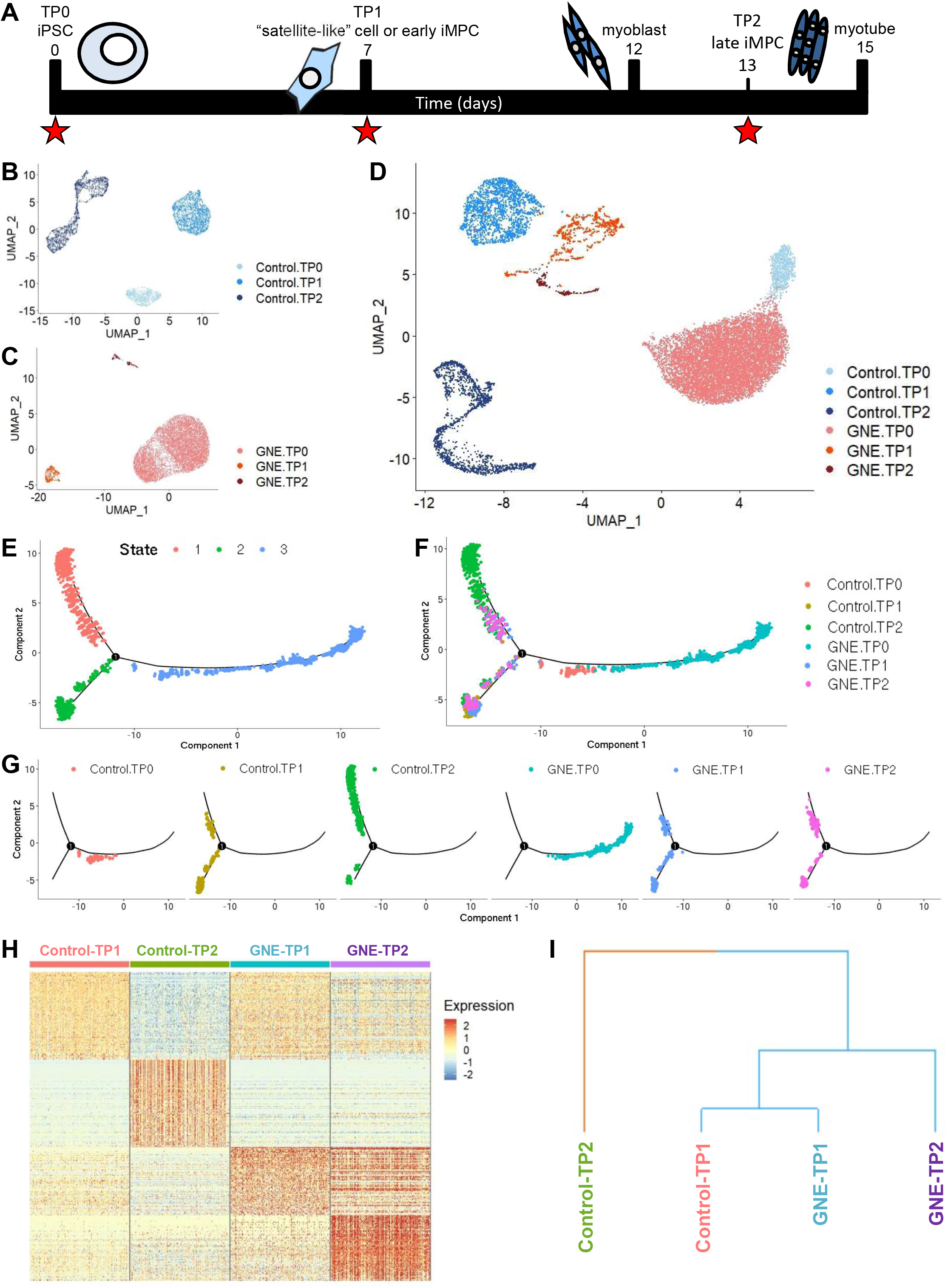
Global transcriptome differences between healthy and GNE patient-derived iPSCs undergoing myogenic differentiation. (A) A schematic depicting the experimental timeline. Red stars correspond to collection of samples for single cell RNA sequencing; first star: Time Point 0; second star: Time Point 1 or early iMPCs; third star: Time Point 2 or late iMPCs. (B-D) UMAP projection of single cell data from (B) control, (C) GNE^1^ (GNE), and (D) control and GNE^1^ combined, containing Time Points 0, 1, and 2. (E-G) Pseudotime trajectory analysis across all time points using single cells from control and GNE^1^ (GNE). (E) The 3 projected state branches were based on differentially expressed genes (DEGs), (F) all single-cell samples mapped onto branches, and (G) individual single cell samples mapped onto branches. (H) Heatmap depicting the log scale of the top 100 DEGs, as represented by 200 cells from each group, for control and GNE^1^ (GNE) Time Points 1 and 2. Red: higher expression and blue: lower expression. (I) Hierarchical clustering depicted via dendrogram using the log scale average of the top 100 DEGs from control and GNE^1^ (GNE) Time Points 1 and 2. Clustering method: complete linkage. Distance measure: correlation.

We then performed a pseudotime trajectory analysis (Monocle) to better capture the state transition defects observed in GNE-TP2 cells. These analyses revealed three distinct cell states (right, upper, lower trajectories), corresponding to the three stages of myogenic differentiation assessed (Fig. 4E). Control-TP0 and GNE-TP0 comprised cell state branch 3 (Fig. 4F - G), the most primitive state prior to induction of myogenesis. The initial branch then split into two divergent branches where branch 2 predominantly contained Control-TP1 and GNE-TP1 cells. These cells also populate a segment of branch 1, closer to the divergence point. Control-TP2 displayed a few cell clusters on the outer edge of the state 2 branch, but are largely distributed along the extended state 1 branch. On the contrary, GNE-TP2 exhibited a trajectory pattern almost identical to the TP1 cells and did not display robust extension into state 1 that was observed in Control-TP2 cells (Fig. 4F-G).

We next visualized the top 100 differentially expressed genes (DEGs) between TP1 and TP2 in a heatmap containing 200 random cells from each of these two time points (Fig. 4H). We found that control and GNE-TP1 samples were most similar whereas Control-TP2 exhibited clear qualitative differences in transcript expression. In contrast to Control-TP2, GNE-TP2 appeared more similar to Control/GNE-TP1. Notably, we observed a cluster of transcripts (dark orange, bottom right) that characterized GNE-TP2, with little to no expression observed in any other sample (Fig. 4H). Hierarchical clustering was performed on the log-normalized average gene expression values from the same 100 DEGs per sample and clearly revealed separation of Control-TP2 from the three other samples, Control-TP1, GNE-TP1, and GNE-TP2 (Fig. 4I).

### Alteration of key myogenic regulatory factor transcripts and the elucidation of a possible contributing mechanism to myogenic dysfunction in GNE iPMCs

Assessment and visualization of myogenic transcript/MRF expression across time points revealed several striking differences between control and GNE samples. First, PAX3 transcript expression at Control-TP1 was 1) detected in the vast majority of cells and 2) expressed at a comparatively high level. At GNE-TP1 there was a reduced number of PAX3-expressing cells and a reduced level of expression per cell (Fig. 5A and C). Second, Control-TP2 exhibited an increase in the number of cells and intensity of expression of MYOD, MYOG, and DES compared to Control-TP1. This effect was not observed in GNE-TP2 samples (Fig. 5A and E-G).

**Figure 5.**
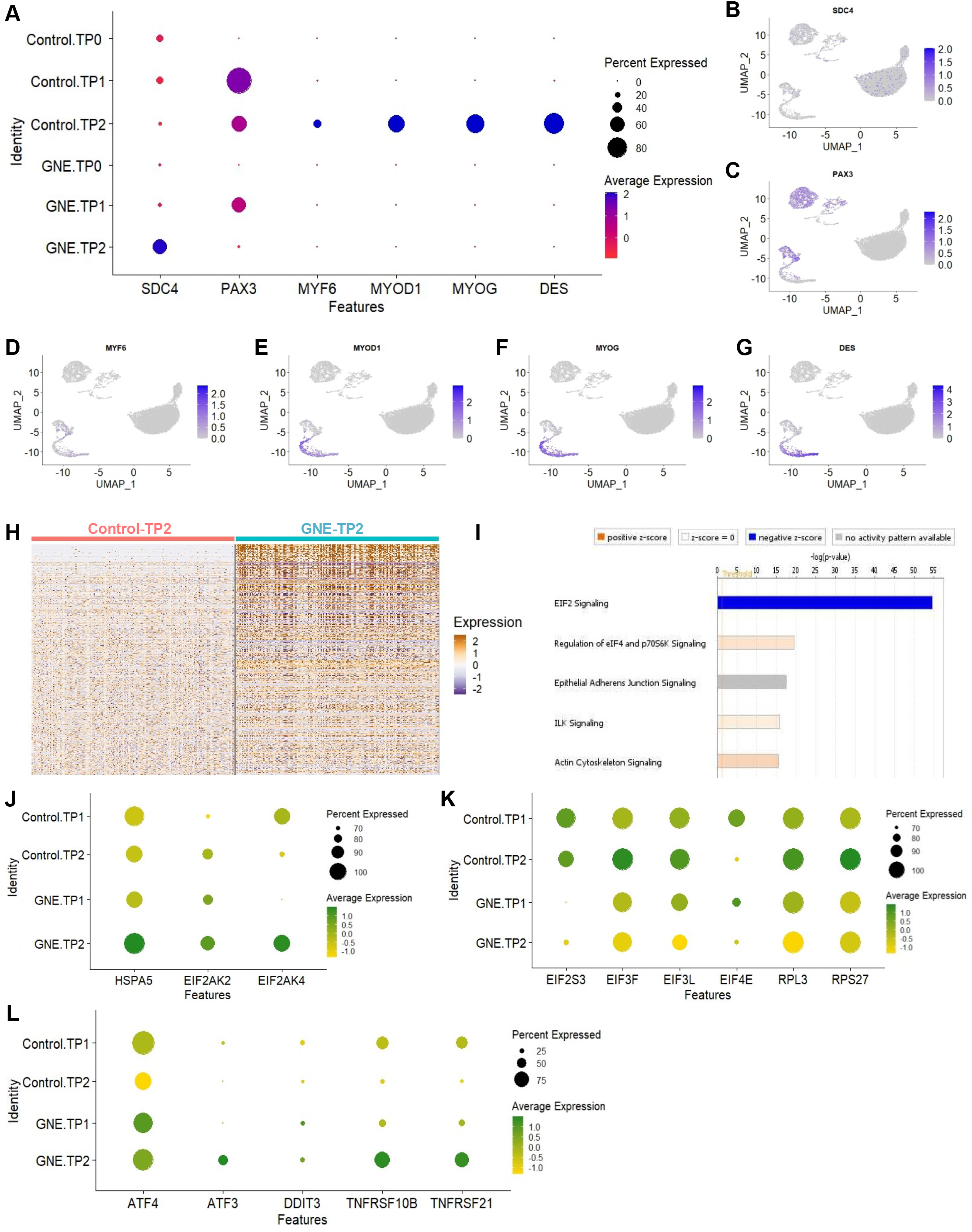
Identification of altered signal transduction pathways in GNE myopathy iMPCs. (A) Dot plots of selected MRFs in Control and GNE^1^ (GNE) samples at Time Points 0, 1, and 2. Blue: higher expression, red: lower expression, dot size indicates proportion of cells expressing specified transcript (%). (B-G) Single cell expression of selected genes from A, depicted via UMAP. Blue: higher expression and gray: lower expression. (H) Heatmap illustrating all DEGs between control and GNE^1^ Time Point 2 (Control-TP2 and GNE-TP2, respectively). Orange: high expression and purple: low expression. (I) Ingenuity pathway analysis (IPA) results using all DEGs found between control and GNE^1^ Time Point 2 (shown in H). Orange: positive z-score, blue: negative z-score, and gray: z-score of 0. (J-L) Selected genes in control and GNE^1^ Time Points 1 and 2, represented by dot plots for involvement (J) upstream of EIF2 signaling, (K) with EIF2 to perform translation, and (L) downstream of EIF2 signaling. Green: higher expression, yellow: lower expression, dot size indicates proportion of cells expressing specified transcript (%).

To address underlying mechanisms that could be contributing to an impaired GNE-TP1 to TP2 transition, we identified DEGs between GNE-TP2 and Control-TP2 iMPCs (Fig. 5H) and performed pathway analyses (Ingenuity). The top five altered biological pathways were: EIF2 Signaling, regulation of eIF4 and 7056K signaling, epithelial adherens junction signaling, ILK signaling, and actin cytoskeleton signaling (Fig. 5I). EIF2 signaling was selected as our top candidate as 1) EIF2 exhibited the highest pathway significance/z-score, 2) EIF2 has been previously implicated in stress granule/aggregate formation (Buchan and Parker, 2009), and 3) EIF signaling is associated with MRF translation (Zismanov et al., 2016). A closer examination of EIF2 signaling DEGs contributing to pathway enrichment revealed several findings. First, we observed progressive upregulation of several upstream factors: (i) Heat Shock Protein Family A Member 5 (HSPA5), which encodes the binding immunoglobulin protein (BiP), a regulator of stress signals from the endoplasmic reticulum (ER), (ii) Protein Kinase R (PRK), encoded by the Eukaryotic Translation Initiation Factor 2 Alpha Kinase 2 (EIF2KA2), which is involved in phosphorylation of the eukaryotic initiation factor 2 subunit 1 (eIF2a) when there is a presence of double stranded RNA, and (iii) the General Control Nonderepressible 2 (GCN2), encoded by EIF2AK4, which is also involved in the phosphorylation of eIF2a, but typically under conditions of amino acid starvation (Proud, 2005). Second, we noted oppositely trending expression of components of the translation pre-initiation complex; increasing expression was observed at the Control-TP1/TP2 transition as opposed to a downwards trend of these same factors at the GNE-TP1/TP2 transition (Fig. 5K). Third, we found evidence of altered EIF2 pathway activity downstream of phosphorylated eIF2a including differential expression of (i) Activating Transcription Factor 4 (ATF4), ATF3, (iii) the DNA Damage Inducible Transcript 3 (DDIT3), (iv) the TNF Receptor Superfamily Member 10B (TNFRSF10B, also known as Death Receptor 5 (DR5)), and (v) TNFRSF21 (also known as DR6) (Fig. 5L). Together, these data provide molecular evidence for GNE-associated myogenesis defects and implicate EIF2 signaling alterations in GNE iMPCs.

## DISCUSSION

This study describes the generation of a novel patient-derived, cell-based GNE model that provides numerous advantages over existing GNE model systems. First, the utilization of two (and potentially more) GNE patient-derived samples of varying pathological severity (Fig. 1) permits modeling of diverse disease severity states and may facilitate genotype-to-phenotype studies not easily performed using mouse models (Fig. 2). Second, as our GNE model is cell-based and patient-derived, there is robust scaling potential for drug screening and/or therapeutic efficacy studies that are labor intensive and technically challenging using animal models. Third, patient-derived samples are relatively easy to obtain, are noninvasive, and reprogramming iPSCs is now a standard and cost-effective platform. Fourth, iPSC-based models permit maintenance of a patient’s entire genetic identity. Gene-by-gene and gene-by-environment interactions are critical when trying to understand the variable expressivity of disease-causative primary mutations and their contribution to clinical phenotype and disease mechanism. For example, inherited myopathies, including GNE myopathy, can manifest with intrafamilial variability in regard to clinical phenotype, age of onset, or disease progression (Dotti et al., 2018; Stober et al., 2010; Diniz et al., 2016). In addition, there is evidence of genetic modifiers, accompanying the primary mutation, modulating the phenotype (Bello et al., 2016; Souza et al., 2020). Therefore, a model system able to capture gene variability while also recapitulating the key pathological characteristics becomes crucial for elucidating yet-to-be discovered contributors to disease etiology.

Although there are many benefits associated with cell-based models, there is one drawback that cannot be overlooked. Extensive and accurate physiological muscle studies, such as muscle force measurements, cannot be conducted using cells, thereby making animal models superior to cell-based models in this regard. In order to overcome this obstacle, there are efforts currently underway to introduce differentiated human iPSCs into mice for *in vivo* physiological assessments. For example, a new model to study Alzheimer’s disease (AD) was created where normal human embryonic stem cells or iPSCs that were derived from a patient with fronto-temporal dementia, were differentiated into neural precursors and implanted into mice to study the etiology of human AD *in vivo* (Espuny-Camacho et al., 2017). Another example from Espuny-Camacho and colleagues found that cortical neurons, differentiated from human embryonic stem cells and implanted into the mouse brain, were able to produce human cerebral cortex-like characteristics *in vivo* (Espuny-Camacho et al., 2013). Such studies highlight the relevance and potential of transplant-based approaches to study human disease in animals. Indeed, we believe that the healthy and GNE patient-derived iMPCs described herein are well-suited for these types of *in vivo* applications.

To our surprise, our single-cell RNA sequencing data revealed little to no expression of MyoD and MyoG transcripts in GNE cells at all-time points (Fig. 5), which on the surface, seems at odds with the immunofluorescence data reported during GNE iPSC differentiation (Fig. 3). One possible explanation, supported by growing literature in the satellite cell field, is that MRFs are subject to substantial post-transcriptional and post-translational modification (Puri and Sartorelli, 2000; Crist et al., 2012; Hausburg et al., 2015; Jin et al., 2016; Asp et al., 2011; Blum et al., 2012; Boutet et al., 2012). For example, Crist and colleagues identified a pathway by which Myf5 mRNA is sequestered to mRNA/protein granules (RNPs) and is translationally “paused” with the help of microRNA-31, in quiescent satellite cells. When satellite cells undergo activation, Myf5 transcript is released for rapid translation, Myf5 protein accumulation occurs, and myogenic differentiation can progress (Crist et al., 2012). In our system, it is possible that *de novo* Myf5 transcription is very low, with most gene regulation occurring post-transcriptionally. Additionally, Myf5 is not the only MRF reported to undergo post-transcriptional regulation. MyoD is a target of multiple RNA binding proteins including HuR (an mRNA stabilizing factor) and Tristetraprolin (TPP, an mRNA decay factor) allowing for precise control of MyoD transcript levels, and thus the timing of MyoD protein induction and satellite cell activation. In one example, TTP is inactivated via p38/MAPK signaling permitting translation and rapid accumulation of MyoD protein (Hausburg et al., 2015). We postulate that in the context of iPSC-based myogenic differentiation, post-transcriptional (and likely post-translational) gene regulation play major roles in MRF protein expression. Future studies, however, are needed to confirm this hypothesis and determine the full extent of MRF gene regulation in differentiating iMPCs.

Despite subtle differences between the mRNA and protein studies, both data sets clearly point to substantial deficiencies in myogenic differentiation. Our immunofluorescence data show altered expression of MRFs (Fig.3) which ultimately culminate in an inability of GNE^1^-derived stage 3 cells to express myosin heavy chain (Fig. 2J, 3J-K). Additionally, MyoG appears to exhibit cytoplasmic localization in some GNE cells, a phenomenon rarely seen in control samples (Fig. 3J). These studies are backed by scRNAseq analysis corroborating myogenesis defects at a global transcriptional level. The discovery of myogenesis/regenerative defects in GNE-derived cells has significant clinical implications. Accumulating evidence in other muscle pathology contexts suggest that targeting muscle regeneration can have a therapeutic benefit. For example, multiple studies investigating age-associated sarcopenia report defects in muscle regeneration (Bernet et al., 2014; Cosgrove et al., 2014). Defective p38α/β signaling in muscle satellite cells appears to be a major driver of this dysfunction and inhibition of the p38/MAPK signaling axis is sufficient to boost the self-renewal of aged stem cells and improve muscle regeneration (Bernet et al., 2014; Cosgrove et al., 2014). Regeneration-targeting interventions are also beneficial in the context of Duchenne muscular dystrophy (DMD). In a DMD mouse model, satellite cell transplantation can alleviate several DMD pathological hallmarks including reductions in maximal isometric tetanic force, specific force, myofiber size/number, and dystrophin expression (Arpke et al., 2013). In a second DMD study, activation of epidermal growth factor receptor (EGFR) and Aurora kinase A (Aurka) in satellite cells enhanced asymmetric satellite cell division, allowing for increased production of cells committed to myogenic differentiation. These DMD mice displayed increases in specific and max tetanic force 30 days post treatment as well as increased tibialis anterior muscle mass, cross-sectional area, and increased number of myofibers (Wang et al., 2019). Lastly, in mouse models of obesity-associated muscle dysfunction, nicotine injection into injured extensor digitorum longus muscles resulted in an increase in the number of myofibers via enhanced muscle regeneration (He et al., 2019). Together, these studies underscore the potential of targeting muscle regeneration to improve muscle histopathology, and possibly, muscle physiology. Our data suggest that this approach could - and should - be explored in the context of GNE myopathy.

## MATERIALS AND METHODS

### Human muscle samples

Muscle biopsies were obtained from patients for diagnostic purpose and conventional histochemical studies were performed on fresh-frozen tissue as previously described (Niu et al., 2018). p62/sequestosome was immunolocalized on 10 μm thick frozen muscle sections with rabbit polyclonal antibody (Abcam) at dilution 1:200 and visualized with secondary antibodies using immunoperoxidase. Use of residual diagnostic muscle biopsy tissues was approved by the Mayo Clinic Institutional Review Board (IRB 13-007054 (MM)); skin biopsies were collected and de-identified as described and approved by Mayo Clinic’s IRB (IRB 15-006983 (MM) & 13-007298 (DW)).

### Patient biopsies and fibroblast isolation

The biopsy site, located on the inner-upper arm, was cleaned with alcohol wipes and local anesthetic was applied. The site was scrubbed with an antimicrobial solution (chloroprep) and a 4mm punch biopsy was obtained and stored in PBS or DMEM with 1% pen/strep. Tissue was washed using DPBS and then minced with 100 μL0.25% trypsin. Tissue was collected in a conical tube and centrifuged for 5 min at 800xg, supernatant aspirated, and suspended in 6 mL fibroblast media. Two T-25 flasks were coated with 0.1% gelatin and the resuspended sample was split between the two flasks and incubated for 14 days at 37°C. After the initial 14 days, media was changed 2 times (2x) per week with 5 mL fibroblast medium. After 7-10 days, the media was aspirated, 2 mL of TrypLE was added, cells incubated at 37°C for 5-10 minutes, and the reaction quenched with 5 mL fibroblast media. Cells were collected into conical tubes, centrifuged for 5 min at 800xg, supernatant aspirated, and cells resuspended in 10 mL fibroblast media. 5 mL cells were plated into T-75 flasks containing 15 mL fibroblast medium, incubated at 37°C with a media change 2x per week. Once cells became 80-90% confluent, they were harvested and plated (as stated above) in T-150 flasks containing 20 mL fibroblast media and fed 2x per week with new media. Reprogramming was performed once cells reached 80-90% confluency.

### iPSC reprogramming

Fibroblasts were reprogrammed using Sendai virus-Cytotune 2.0 as described by manufacturer following the Reprogram fibroblasts Feeder-Free instructions. Briefly, fibroblasts were transduced using the CytoTune 2.0 Sendai reprogramming vectors and incubated at 37°C overnight. The next day, media was changed to new fibroblast media and then for 6 days, fibroblast medium was changed every other day. Vitronectin was used to coat plates and the fibroblasts were harvested and plated at 2×10^4^−1×10^5^ and incubated overnight. Media was then changed to complete Essential8 Medium (Thermo) and changed every day for the next ~20 days. Cultures were monitored for iPSC colonies which were picked and cultured using new vitronectin-coated dishes for expansion.

### iPSC culture and quality control

iPSCs were maintained on Geltrex (Thermo Fisher)-coated plates in mTeSR1 (STEMCELL Technologies). Passaging occurred every 3-4 days using ReLeSR (STEMCELL Technologies) and DPBS 1X (Gibco) for washing. Germ layer differentiation was performed essentially as described by manufacturer. iPSCs were plated at a density of 200,000 cells/cm^2^ for ectoderm and endoderm differentiation and 100,000 cells/cm^2^ for mesoderm. Ectoderm Medium (STEMCELL Technologies) with 10 μM Y-27632 (STEMCELL Technologies) was used as plating medium for cells undergoing ectoderm differentiation. For cells undergoing mesoderm or endoderm differentiation, the plating media used included mTESR1 with 10 μM Y-27632. After 24 hours, plating mediums were changed to ectoderm, mesoderm, or endoderm mediums (STEMCELL Technologies) respectively. Media change occurred every day for 5 days for mesoderm and endoderm and 7 days for ectoderm.

### iPSC/iMPC myogenic differentiation

Skeletal muscle differentiation occurred in a three stage process essentially as described, briefly: (1) iPSCs were differentiated to myogenic precursors by dissociating iPSCs with Accutase (Sigma) for 5 minutes, resuspended in Skeletal Muscle Induction Media (Stage 1 media) (Genea Biocells), centrifuged at 1200 rpm for 2 minutes, decanted, and suspended in Stage 1 media. Cells were counted and plated in BioCoat collagen I 24-well plates (Corning) at 7,500 cells/cm^2^. Stage 1 media was changed every 2 days until cells reached confluence, (2) myogenic precursor cells were further differentiated to myoblasts by dissociation of myogenic precursor cells using 0.05% Trypsin-EDTA for 5 minutes, trypsin was neutralized using 10% FBS (Gibco) in DMEM (Gibco), cells centrifuged for 4 minutes at 1200 rpm, and resuspended in Skeletal Myoblast Medium (Stage 2 media) (Genea Biocells). Cells were counted and plated in 24 well Collagen I coated plates at 10,000 cells/cm^2^. Stage 2 media was changed every 2 days until cells reached confluency. (3) Myoblasts were differentiated to myotubes by changing Stage 2 media to Skeletal Myoblast Medium (Stage 3 media) (Genea Biocells). Stage 3 media was changed every 2 days until the formation of myotubes.

### Sanger sequencing/mutation confirmation

iPSC samples were used for DNA isolation using the DNeasy Blood and Tissue Kit (Qiagen). PCR was performed using PCR SuperMix (Thermo) following manufacturer’s instructions. All PCR products were enzymatically cleaned using ExoSap-IT (Thermo). Sequencing primers were added and samples sent to GeneWiz for Sanger sequencing. 4Peaks software was used for trace visualization and presence of mutation verification.

Primers were purchased from Integrated DNA Technology and sequences used are as follows (Forward/Reverse/Sequencing Primer):

GNEc.479G>A: 5’-CCAGGCTACACACAATTGTGAGGGG-3’/5’-GAAGTTTGTCATAGGAAGGGCAGCC-3’/5’-GCTGCCAGATGTCCTTAATCGCCTG-3’

GNEc.1835G>T: 5’-GATGCCTAGTGGGCTTCAGCTGTC-3’/5’-CCACCACCACCCCTGGGGAGG-3’/5’-CCCTCGCTGTAGGAATCGGTGGTG G-3’

GNEc.2179G>A and c.2218G>A: 5’-GCCTCCCACTGCATGCCGTGG-3’/5’-GTGGGGAAAGGTGACTCTGGAAGAGG-3’/5’-CCCTCCCTTGTGATCCTCTCCGG-3’.

### Immunofluorescence

Cultured cells from the indicated time points were washed with PBS and fixed in 24 well plates using 4% Paraformaldehyde Solution (Santa Cruz) for 20 minutes at room temperature, washed 2x with PBS, and the stored in PBS at 4°C until ready for use. Cells were permeabilized using 0.2% Triton X-100 (Fisher) in 1:10 dilution Blocking Buffer (Dako) for 30 minutes at room temperature. Cells were washed for 5 minutes using 1X Wash Buffer and then blocked by adding 8 drops of Protein Block Serum-Free (Agilent). Primary antibodies were diluted in diluent (Dako) and incubated at room temperature for 1 hour, washed 2x with Wash Buffer for 5 minutes, and incubated with secondary antibodies in diluent for 30 minutes at room temperature. Cells were washed 2x with Wash Buffer for 5 minutes, incubated with Nunc Blue (Invitrogen −2 drops/mL in PBS) for 5 minutes, washed with PBS, and stored in PBS at 4°C for up to 2 weeks for imaging. See antibody details in Supplemental Table 2. Images were taken with BioTek Cytation5 imaging reader using Gen5 (v3.06) software.

### Image and statistical analyses

Cell Profiler (version 3.1.9) (McQuin et al., 2018) was used to quantify nuclei positive cells by setting the diameter and upper/lower thresholds in control images and applying the same settings to the GNE images, except for the stage 3 quantifications in Figure 3K-L which was performed manually. Each n is the average percent positive nuclei for 5 images per well. GraphPad Prism 8 (version 8.4.2) was used for all statistical analyses. One-way ANOVAs were performed with Bonferroni Multiple Comparisons (BMC) where significance was p < 0.05. All numerical data is reported as mean ± s.e.m.

### Single-cell mRNA sequencing

Three wells, from a 24 well plate (Corning), of cells were scraped and combined from the indicated time points and suspended in DPBS. Cells were then assessed for viability and count using Vi CELL XR (Beckman Coulter) analysis version 2.42. Samples with the highest cell count and viability were selected for further single cell RNA sequencing processing. 10X Genomics single cell reagent 3’ v2 was used to prepare the cDNA libraries and then were sequenced on the Illumina HiSeq 4000, with paired-end 100bp reads. 10X Genomics Cell Ranger Single Cell Software Suite (v2.2.0) was used to demultiplex raw base call (BCL) files generated from the sequencer into FASTQ files (Cellranger mkfastq command) and to perform alignment to the hg38 genome, filtering, barcode counting and UMI counting (Cellranger count command). The gene expression matrices files were used for the subsequent analyses. Seurat version 3.2.0 (Stuart et al., 2019) (https://github.com/satijalab/seurat/releases/tag/v3.2.0) was used in RStudio (http://www.rstudio.com/. Version 1.3.959. Release Name: Middlemist Red) using R Version 4.0.2 (https://cran.r-project.org/bin/windows/base/old/4.0.2/) to analyze data and filter out cells that were poor quality. Coding was performed predominately following the Satija lab Seurat tutorial (https://satijalab.org/seurat/v3.1/pbmc3k_tutorial.html). Each sample was assessed individually and threshold values were assigned for low unique molecular identifiers (threshold 1,800 - 10,000), low gene detection (threshold 900 - 3,000), and high mitochondrial transcript fraction (threshold 10-25%). All data was i) normalized using NormalizeData, ii) scaled using ScaleData, iii) run with principal component analysis (PCA) using the RunPCA command, and iv) UMAP using RunUMAP.

Pseudotime trajectory was performed using Seurat version 3.2.0 (Stuart et al., 2019) (https://github.com/satijalab/seurat/releases/tag/v3.2.0) in RStudio (http://www.rstudio.com/. Version 1.2.5042.1. Release Name: Double Marigold), R Version 3.6.2 (https://cran.r-project.org/bin/windows/base/old/3.6.2/), with monocle version 2.14.0 (Qiu et al., 2017b; Qiu et al., 2017a; Trapnell et al., 2014) essentially as described (http://cole-trapnell-lab.github.io/monocle-release/docs/#constructing-single-cell-trajectories).

Hierarchical clustering was performed by TIBCO Spotfire Analyst 7.11.2.4 using the average log scale gene expression for each time point for the top 100 differentially expressed genes as found by the R command FindAllMarkers, with expression in a minimum of 25% cells and greater than a log fold-change of 0.25.

## Acknowledgements

We are grateful for Dr. Dennis Wigle for the use of the control iPSC line in this study. We are also indebted to Dr. Nathan Staff and his team for facilitating the iPSC reprogramming. Finally, we would like to thank the Mayo Clinic Genome Analysis Core for performing the single-cell RNA sequencing and genome alignment.

## Data availability

Data from this study is available in the Sequence Read Archive (accession number).

